# Minimal mean-field gated parietal circuit model for flexible perceptual decisions

**DOI:** 10.1101/2025.09.09.675139

**Authors:** Brendan Lenfesty, Amin Azimi, Saugat Bhattacharyya, S. Shushruth, KongFatt Wong-Lin

## Abstract

Flexible perceptual decision-making requires rapid, context-dependent adjustments, yet the neural circuit mechanisms underlying its parsimonious representations remain unclear. Here, we propose a minimal mean-field neural circuit model that integrates sensory evidence and selects actions via distributed neuronal encoding, guided by data from a task that dissociates perceptual choice from motor response – abstract perceptual decision-making. The model’s nonlinear gating of action selective (AS) neurons replicates parietal cortical activity observed during task performance. Critically, recurrent excitation within the evidence integration (EI) population supports sensory evidence accumulation, working memory for sequential sampling, and reward rate optimisation. Moreover, the dynamics of EI and AS neuronal activities in the same model respectively mirror parietal neuronal activities related to sensory evidence encoding and ramping-to-threshold firing in a separate reaction-time task, while suggesting that decision readout engages both neuronal populations. The model also predicts decision interference in a novel two-stage decision version of the task, accounting for choice accuracy decrements observed in other experiments while predicting slower decisions. Together, these findings propose a minimal mean-field circuit-level mechanism unifying perceptual, memory-based, and abstract decision-making.

## Introduction

Perceptual decision-making refers to the transformation of sensory input into a unitary decision (Hanks & Summerfield, 2017; O’Connell et al., 2018). Traditionally, this has been attributed to the temporal integration of sensory evidence — a process where information is integrated over time before a decision commitment (Gold & Shadlen, 2007). This framework has been supported by neural and behavioural data, and computational modelling (Wang, 2008; O’Connell et al., 2018). However, many classical task paradigms (e.g. Shadlen & Newsome, 2001; Roitman & Shadlen, 2002) conflate sensory evidence integration with motor action selection, making it difficult to parse their individual contributions.

Recent studies have begun disentangling these components by exploring stimulus-action mappings that identify more abstract neural signals agnostic to sensory or motor modalities (O’Connell et al., 2012; Twomey et al., 2016; Ede & Nobre, 2024; Gherman et al., 2024; Xie et al., 2024). These findings suggest that abstract neural representations support flexible decision-making across changing contexts, even in the absence of immediate action cues (Okazawa & Kiani, 2023) (Figure 1A). Animal studies complement this view (Wang et al., 2019; Shushruth et al., 2022; Charlton & Goris, 2024; Hasnain et al., 2025). Notably, Shushruth et al. (2022) showed that there were neurons in the lateral intraparietal cortical area (area LIP) in monkeys which were quiescent during sensory stimulus presentation but began integrating motion evidence only after a response cue was presented — indicating memory-based, action-gated decision encoding. Complementing this, Zylberberg & Shadlen (2025) identified distinct neuronal subpopulations in area LIP involved in evidence integration, choice, and decision uncertainty. This highlights the heterogeneity of area LIP, but it remains unclear how a neural circuit can flexibly and rapidly adapt to perform different tasks without (re-)learning.

**Figure 1.**
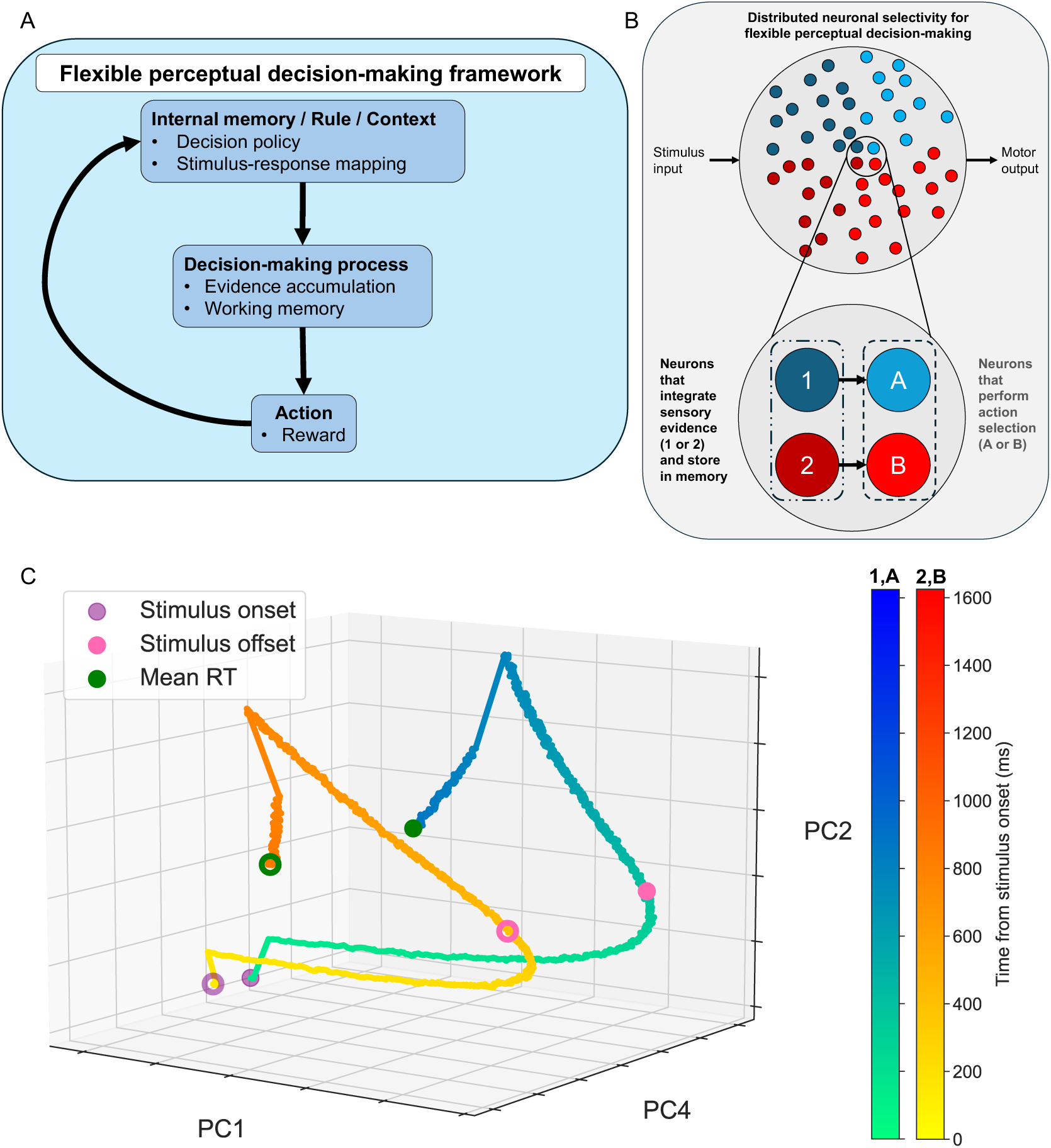
Flexible perceptual decision-making framework and a parsimonious neurocomputational model. **(A)** Flexible perceptual decision-making follows rules or context, guided by internal memory, to determine decision-making process and action selection. Action outcome in turn updates memory and rule (Okazawa & Kiani, 2023) **(B)** Top: Neurons with various selectivity or tuning, as in some reservoir network models (illustrated by different colour labelling). Bottom: Parsimonious representation with neuronal populations selective to sensory stimulus of type 1 or 2 (e.g. leftward or rightward visual motion direction), or selective of action for choice target A or B (e.g. blue or yellow coloured choice targets). For simplicity, we ignore spatial selectivity. **(C)** Minimal mean-field model’s neural trajectories for two different choices projected onto PCA state space can exhibit low-dimensional unfolding over time along curved manifold. Model discriminated an ambiguous stimulus with fixed time duration of 466 ms (Methods). Trial-averaged activities for two different selected choices. Coloured circles: Different time stamps within a trial.

Recent computational models of decision-making often use abstract forms of recurrent neural networks, particularly reservoir network models to understand flexible decision dynamics (Sussillo & Barak, 2013; Song et al., 2017; Xu et al., 2024; Langdon & Engel, 2025; Genkin et al., 2025) (Figure 1B, top). A prominent framework is the “neural manifold”, which posits that neural activity during decision-making evolves along low-dimensional trajectories embedded within high-dimensional neural activity spaces (Langdon et al., 2023; Perich et al., 2025), extending classic decision models (Bogacz et al., 2006; Wong & Wang, 2006). Further, recent computational work shows that distinct tasks can be represented in orthogonal subspaces of this manifold, allowing task flexibility (Liu & Wang, 2024). However, many models utilise high-dimensional random recurrent circuit connectivity comprising abstract neurons, which do not directly provide biological interpretability or mechanistic explainability. Critically, such modelling approaches typically require learning model parameters for every task under investigation. Consequently, it remains unclear whether these trained models can be directly applied to a different, previously unseen task. Moreover, it is uncertain whether successful transfer to a different task is possible using only a minimal mean-field model that can be directly linked to experimentally observed neuronal firing rate activities.

To address this, we propose a biologically plausible minimal mean-field computational model comprising parietal cortical neuronal populations and reverse engineer functional selective populations to separately encode evidence integration and gated action selection (Figure 1B, bottom). This neural circuit model integrates the principles of nonlinear dynamics, memory fidelity and gain modulation, providing a unified foundational framework for perceptual, memory-based, and abstract decisions, that mechanistically explains and predicts how flexible decisions are computed across diverse perceptual decision-making task contexts.

## Methods

### Experimental task

The mean-field model to be developed was based on our previous experimental study (Shushruth et al., 2022). The study involved designing a two-alternative forced-choice experiment for two monkeys (AN and SM), with slight variation, to dissociate sensory evaluations from motor actions – abstract decision-making. The task uses random-dot motion kinematogram stimulus to visually discriminate coherent motion direction while transforming the perceptual decision into choosing motion associated coloured targets. To choose correctly, blue (yellow) coloured target was associated with rightward (leftward) coherent motion for monkey AN but the opposite for monkey SM. After motion stimulus offset, a fixed delay period (Go-task = 333 ms and Wait-task = 200ms) ensured that sensory processing and motor planning were temporally unyoked. Following the delay, coloured choice targets appeared in random positions on the screen. Monkey AN then responded immediately (Go-task) while monkey SM waited until a fixation point was extinguished before being cued to indicate their choice (Wait-task). For further details on the original experiment, please refer to Shushruth et al. (2022).

### Behavioural data analysis

Choice accuracy was assessed for each monkey as the proportion of correct trials at each motion coherence level and combining all sessions with valid responses. For monkey AN, go-reaction times (go-RTs) were also analysed across correct and error trials, combined for both choice targets. This provided a behavioural index of how task difficulty influenced both response accuracy and speed.

### Neuronal data preprocessing and epoch grouping

To investigate motion coherence responses during decision formation, we re-analysed single-unit recordings in area LIP from Shushruth et al. (2022). Spike trains were smoothed using a zero-phase 40 ms boxcar filter, a small enough window to smooth but still capture task related activity. Neuronal firing rates were then standardised by dividing each trial’s activity by the maximum rate observed within its session. Trials were grouped by motion coherences, namely, zero (0%), low (4% & 8%), medium (16%), and high (32% & 64%) coherence, and aligned to three key epochs: stimulus onset, target onset, and saccade.

### Neuronal selection

Given the heterogenous choice encoding of LIP neurons, we hypothesised that neurons with stronger relationship to the decision contributed more strongly to downstream decision process (Supplementary Figure S1). Hence, we searched for differential firing rate activities based on difficulty during each session. Specifically, we identified decision-relevant neuronal responses to motion coherence by analysing the trial-averaged LIP neuronal firing rates from each recording session (Supplementary Figures S2 and S3) across task difficulties during the target-onset epoch.

For monkey SM, this revealed one neuron (neuron 25) with the clearest (separable) encoding of motion coherence after target onset; we did not see a similar session for monkey AN (Supplementary Figures S2 and S3). A quantitative selection criterion was utilised, with correlation between each neuron (sessions) computed, summated and averaged across task difficulties (Supplementary Figures S4, S5, S6 and S7). This led to the inclusion of 20 more neurons that showed a Spearman’s correlation greater than 0.5 with respect to neuron 25. No coherence-based selectivity was observed for monkey AN using this method, and hence all (n = 28) recorded neurons were retained.

### Statistical analysis

For both monkeys, we examined trial-wise firing rate differences across coherence conditions during the target-onset epoch. Pairwise comparisons (e.g., zero vs. low coherence) were tested using the Mann-Whitney U test at each time point post-target onset. Multiple comparisons across time were corrected using the Benjamini-Hochberg false discovery rate procedure. Time points showing significant difference between coherences (p < 0.05) were marked in Figure 2D. This figure reflects combined firing rates from all 28 neurons in AN and 21 neurons in SM, aligned to trials where the chosen target appeared in the neuron’s response field.

**Figure 2.**
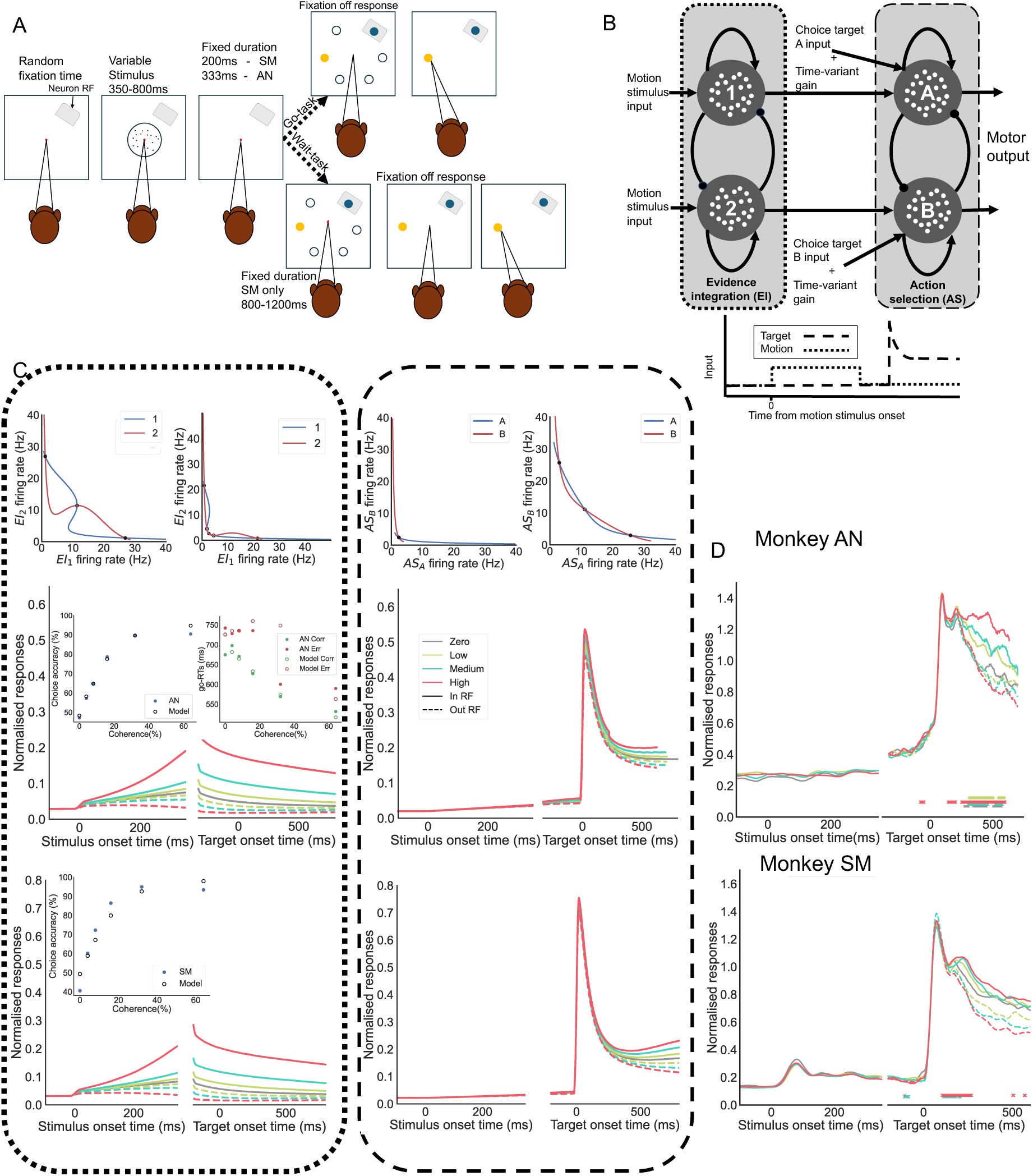
Model recapitulates neuronal and behavioural observations in an abstract perceptual decision task. **(A)** Experimental task (Shushruth et al., 2022) of perceptual decisions abstracted from action selection. Monkey AN discriminated motion coherent direction and made a saccade to blue (yellow) target for perceived rightward (leftward) motion, this was reversed for monkey SM. Targets might appear in or out of neuronal response field (grey patch). Go-task: Upon target onset, monkey AN was free to saccade with noted reaction time (RT) from target onset (go-RT). Wait-task: Monkey SM waited for cue to saccade. **(B)** Neural circuit model with neural group (dotted) that performs sensory evidence integration (EI) for different motion coherences and stores decision in delay period, while sending information to action selection (AS) group (dashed). Time-variant gain parameters varied between monkeys. Bottom: input protocol. **(C-D)** Model fit to behavioural data from monkeys AN (top) and SM (bottom). **(C)** The model captured LIP neural activity across different motion coherences (dotted region) via AS neural dynamics (dashed region), and additionally predicted the activity of unobserved EI populations (dotted region). Inset: EI phase planes during stimulus presentation with zero motion coherence and the delay period (left and right, dotted); AS phase planes are shown before and after target onset (left and right, dashed). **(D)** Corresponding LIP neuronal activity from monkeys. Reanalysis of LIP neurons revealed significant correlations with the animals’ choices (Mann–Whitney U test, corrected using the Benjamini–Hochberg false discovery rate procedure, p < 0.05). Crosses: significant time points.

### Model architecture

To investigate the neural mechanisms underlying temporally flexible decision-making, we extended the neurobiological mean-field recurrent network model of Wong and Wang (2006), which consists of two effectively mutually inhibited neuronal populations governed by slow NMDA-mediated synaptic dynamics. In particular, we created two replicas of the model and coupled them with unidirectional connectivity. One module comprised sensory evidence integration (EI) to integrate for forming and storing perceptual decision, while the second module transforms EI output as input for action selection (AS) (Figure 1B, bottom). As we were considering only two choice tasks, each module comprised two competing neural populations, labelled as sensory (stimulus) based decisions “1” and “2” in EI, and actions “A” and “B” in AS. Each neural population’s activity is governed by recurrent network dynamics, with nonlinearity in the neuronal input-output functions and NMDA-mediated synaptic currents (see below).

For simplicity, we assumed the EI neural group to continuously feed its output as input to the AS neural group such that neural population 1 in EI was connected to AS neural population A (Figure 1B, bottom). Similarly, neural population 2 was connected to AS neural population B (Figure 1B, bottom). Moreover, each neural population comprised self-excitatory connectivity representing neuronal interactions within each population. When a stimulus, a mixture of sensory information 1 and 2, is presented, the network model could simulate evidence integration and form a perceptual decision (1 or 2). Due to strong recurrent reverberation, this decision information is stored over time in the absence of stimulus during delay period, before an action (A or B) is selected upon cue (Methods) (Roitman & Shadlen, 2002; Shadlen & Newsome, 2001; Shushruth et al., 2022).

### Neuronal input-output function

The neuronal population’s nonlinear input-output functions, 𝑟_j_, follow Wong and Wang (2006) and Niyogi and Wong-Lin (2013):

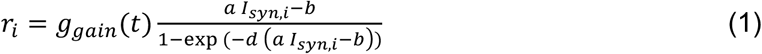

where 𝑖 denotes the 𝐸𝐼_1_, 𝐸𝐼_2_, 𝐴𝑆_𝐴_ or 𝐴𝑆_𝐵_ neural population, and parameters a, b and d were based on derivation via spiking neuronal model (Wong & Wang (2006)). The time-varying gain factor *𝑔_gain_*(𝑡) was 1 for EI populations while for AS populations was described by (Niyogi & Wong-Lin (2013)):

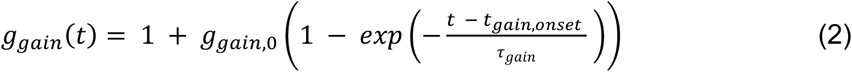

which was turned on at 𝑡_*gain,𝑜𝑛𝑠𝑒𝑡*_ whenever stimulus or target was turned on, whichever came first. The gain allowed slow buildup due to urgency, attentional readiness, or anticipatory signal.

Equation (1)’s 𝐼_𝑠𝑦𝑛,𝑖_ was the total synaptic input to the neural population 𝑖. For EI populations 1 and 2, they were described respectively by:

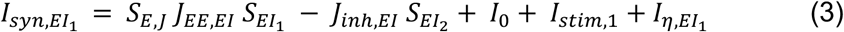

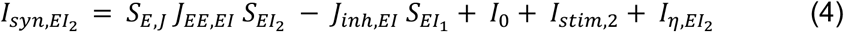

where 𝐼_0_ was biased current, 𝐽_𝐸𝐸_ was self-excitatory coupling strength with multiplicative factor 𝑆_𝐸,𝐽_, and 𝐽_𝑖𝑛ℎ_ the mutual inhibitory coupling factor. The motion stimulus inputs 𝐼_𝑠𝑡𝑖𝑚_ in Equations (3-4) to each of the two EI populations were:

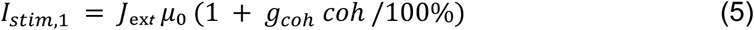

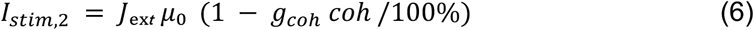

with 𝐽𝖾ₓₜ and 𝜇₀ as the external input coupling strength and parameter for baseline stimulus input (MT neuronal output), and 𝑔_𝑐𝑜ℎ_ the motion stimulus’ gain factor.

The noisy input 𝐼_𝜂_ in Equations (3-4) has motion stimulus gain factor, 𝑔_𝑐𝑜ℎ_, which was 1 for AS populations and the prescribed parameter value for EI populations. 𝐼_𝜂_ was generated via Ornstein-Uhlenbeck process as follows (Wong & Wang (2006)):

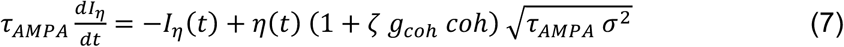

where 𝜏_𝐴𝑀𝑃𝐴_ was the decay time constant of AMPA-mediated synapses, 𝑐𝑜ℎ the motion coherence level, 𝜁 a stimulus-dependent multiplicative noise, 𝜎 the overall noise amplitude, and 𝜂 a random variable that follows a Gaussian distribution with zero mean and standard deviation of one.

The 𝑆_𝐸𝐼1_ and 𝑆_𝐸𝐼2_ in Equations (3-4) were the two populations’ slow NMDA-mediated synaptic gating variables, which were the main dynamical variables in the model, and described by:

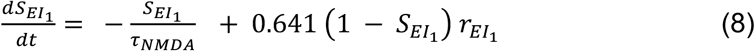

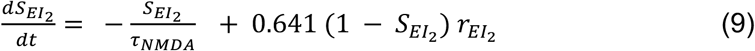

where 𝜏_𝑁𝑀𝐷𝐴_ was the decay time constant of NMDA-mediated synapses.

The synaptic inputs to the AS populations were described by:

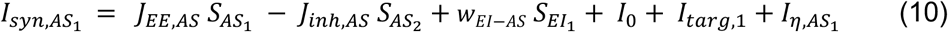

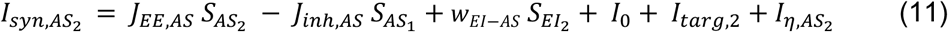

with similar notation meaning as Equations (3-4). The only differences were the feedforward excitatory inputs from corresponding EI populations via the EI-to-AS coupling strength 𝑤_𝐸𝐼–𝐴𝑆_and that instead of the motion stimulus input, we now had the choice target inputs, as described by:

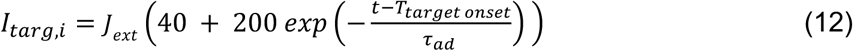

with an initial large spike mimicking change detection, followed by adaptation with time constant, 𝜏_𝑎𝑑_ (Wong et al., 2007). The choice targets, together with the network’s emergent nonlinear dynamics, acted as a gating mechanism for response initiation. Moreover, the AS populations’ dynamics were described in a similar way as in Equations (8-9) and their noise inputs as in Equation (7).

The mean-field model was first fitted to the psychometric (choice accuracy) and chronometric (reaction time) of monkey AN. To allow direct transfer of the model across tasks, the model parameters were maintained in subsequent investigations, except for the gain decay time constant for monkey SM and the first decision threshold for the cognitive interference task (Supplementary Table S1).

### Projection onto neural subspace

To obtain neural manifold from sensory evidence integration to action selection, we projected its trial-averaged neural trajectories onto neural subspace by performing principal component analysis (PCA) on the combination of the average of competing model populations (neural population 1 and population A; and population 2 and population B) for zero coherence to identify any prominent features across trials. First, we applied a 40 ms zero phase boxcar filter to smooth data. The data is then combined for all neuronal populations and fitted with PCA. We then project populations for each of the two choices back into the subspace using the loading weights. For visualisation, we found that the first two and the fourth principal components (PCs), PC1, PC2 and PC4, offered the clearest neural trajectory separability based on the two different choices made at mean reaction time for zero coherence as well as displaying orthogonal subspace when integrating sensory information as compared to when overtly reporting choices via motor response.

### Model simulations and analyses

The code for the analysis and modelling were written in Python Jupyter Notebook. We simulated our model’s stochastic dynamical equations using the forward Euler-Maruyama integration scheme with timestep of 0.5 ms; smaller time step did not affect the results. The model parameter values are summarised in Supplementary Table S1. After initial fitting to Shushruth et al. (2022) the model was simulated and compared across two additional experiments, one which was an actual experiment (Steinemann et al., 2024; Zylberberg & Shadlen, 2025) and another was a thought experiment inspired by cognitive interference studies, e.g. by Wang & Busemeyer (2016). Phase-plane analysis using Python code, was performed by setting the dynamical equations to be zero, and solving the equations algebraically to obtain its nullclines.

## Results

### A minimal mean-field model that can manifest neural manifold during sensory evidence accumulation and action selection

With growing experimental and theoretical evidence of distributed encoding of neurons in perceptual decision-making (Figure 1B, top) (Bollimunta & Ditterich, 2012; Sussillo & Barak, 2013; Mante et al., 2013; Brody & Hanks, 2016; Cueva et al., 2020; Shenoy & Kao, 2021; Steinemann et al., 2024; Zylberberg & Shadlen, 2025; Langdon & Engel, 2025; Asadpour & Wong-Lin, 2025), we reverse engineered and developed a parsimonious and tractable neural circuit model that captured the key functions in perceptual decision-making: sensory evidence integration (EI) and action selection (AS), with neuronal pools selective to one of these functions (Figure 1B, bottom; Methods).

To demonstrate that this mean-field model, although structurally minimal, can still relate to neural manifold known in the literature, we first simulated a generic a two-alternative forced choice task in which the model discriminated ambiguous sensory stimulus with fixed duration before selecting a choice target. By projecting the trial-averaged network’s neural trajectories in principal component analysis (PCA) space (Methods), each representing one of the two choices made, we could observe it unfolding from stimulus target onset till time to respond (Figure 1C). This is akin to observations in multi-neuronal/unit recordings and reservoir network modelling (Sussillo & Barak, 2013; Aoi et al., 2020; Finkelstein et al., 2021; Okazawa et al., 2021; Ashwood et al., 2022; Thura et al., 2022; Genkin et al., 2025). However, the minimal mean-field model offers the simplest instantiation and interpretability of neural manifold in higher dimensional state space.

### Gated mean-field model recapitulates observed abstract decision behaviours and LIP neuronal activity patterns

To constrain the mean-field model, we fitted it manually to experimental data of non-human primates making perceptual decisions abstracted from action plans (Shushruth et al., 2022) (Methods). Figure 2A depicts this experiment in which two monkeys discriminated leftward or rightward on coherent direction of a presented random-dot motion stimulus. After a brief delay period following motion stimulus offset, two coloured choice targets (yellow and blue) were presented. The monkeys reported their decision based on the learned association between direction of motion coherence and colours of choice; choosing yellow (blue) target if leftward motion was perceived and blue (yellow) target if rightward motion was perceived for monkey AN (monkey SM). Monkey AN was free to saccade to one of two coloured choice targets upon their onset and the time taken for the monkey to choose a target was defined as the go-reaction time (go-RT) (Figure 2A). Differing from this free response task, monkey SM instead waited for a cue (fixation offset) to saccade to the appropriate coloured choice target, hence no go-RT was determined (Figure 2A).

We manually fitted the mean-field model’s output to the choice accuracy and go-RT by identifying an appropriate EI-to-AS connectivity while also including an urgency input signal to the AS neural group (Niyogi & Wong-Lin, 2013) (Figure 2B) (Methods). Sensory stimuli 1 and 2 were now represented as leftward or rightward direction of the motion coherence, respectively. Similarly, actions A and B represented choosing blue or yellow coloured choice target, respectively. Otherwise, the model was essentially two connected replicas of the Wong and Wang (2006) model with target inputs similar to Wong et al., (2007) (Methods). Further, as the two monkeys performed slight variation of the task, we varied only the gain time constant such that it was relatively faster for monkey AN due to it undertaking a more urgent (free response) task than that of monkey SM.

After model fitting (Figure 2C, left, inset), the EI neuronal populations could sequentially sample sensory evidence during motion viewing and cache it for later reporting (Figure 2C, dotted region). This was enabled via winner-take-all network behaviour as validated by EI’s dynamical phase-plane analysis (Figure 2C, inset in dotted region). Winner-take-all behaviour during sensory evidence integration was instantiated by two competing fixed-point attractors (Figure 2C, left inset in dotted region) before storing the decision information over time as persistent activity in stimulus absence via multistable working memory attractors (Figure 2C, right inset in dotted region).

Further, although EI’s output was continuously fed as input to the AS neuronal populations, the latter were not substantially activated until gated by the choice target input (Figures 2B-C). This gating mechanism was achieved through the interplay between target input and nonlinear network dynamics via crossing a dynamical (saddle-node) bifurcation point (Wong & Wang, 2006) (Figure 2C, inset in dashed region). Once this “gate was opened”, triggered by choice target inputs, AS neuronal populations reflected graded coherence-dependent decision evidence of EI’s output (compare activities in Figure 2C’s dotted and dashed regions). It should be noted that strong persistent EI activities encoding the committed decision was unnecessary given the brief delay (Figure 2C, right of dotted region). More importantly, the decaying EI activities allowed graded encoding of coherence as input to AS.

To compare the model’s neural activities with LIP neuronal activities, we re-analysed the neuronal spiking activities in area LIP in Shushruth et al. (2022). Due to LIP’s heterogeneous neuronal activity patterns (Supplementary Figures S2 and S3), we selected LIP neurons that exhibited clear separation of activities with respect to motion coherence levels (Methods). We hypothesise that neurons which are more strongly related to decision information would more likely contribute more to the decision outcome (via downstream readout) (Supplementary Figure S1). Our analysis began by inspecting each neuron with the clearest activity separation and other neurons that strongly correlated with it across coherences (Supplementary Figures S4-S7) (Methods). To simplify the analysis, we used trials with the selected blue or yellow targets that were within the response fields of the recorded LIP neurons.

Figure 2D shows the re-analysed trial- and population-averaged LIP activity timecourses. The LIP neuronal firing rate response to motion coherence following target onset suggested target-gated delayed evidence integration, consistent with that of the model’s AS populations (Figure 2B, dashed region). Statistical analysis of neuronal responses on motion coherence validated the differential firing rates during target onset epoch. Crucially these dynamics also suggested the maintenance of motion coherence related information in memory throughout the delay period, as in our model’s EI neurons, but unobserved in the limited experimental recordings. Additionally, we also noticed the slightly elevated LIP neuronal activities prior to target onset, like the model’s AS neural activities due to time-variant gain modulation and input from the EI group. However, in the model, such elevated activities were limited due to the constraint of a low-lying spontaneous attractor state of the network (Figure 2C, left inset in dashed region) (Wong & Wang, 2006).

### Model suggests monkey attained optimal reward rate with intermediate recurrent excitatory strength

The graded AS activities were derived from the graded stored activities of the EI group across the delay period due to its activities decaying over time (Figure 2C). Hence, we next investigated how different memory strengths in the model could affect decision formation and storage, and subsequently decision behaviours. This was achieved by varying a multiplicative factor 𝑆_𝐸,𝐽_ of the self-excitatory connectivity (Methods) of the EI neuronal populations from its control value of 𝑆_𝐸,𝐽_ = 0.975.

When the EI’s self-excitatory connectivity was too strong (𝑆_𝐸,𝐽_ =1.25), EI neurons became highly excitable and the ‘losing’ (out of response field) EI neuronal population’s activity (Figure 3A, left, dashed) was also elevated for all motion coherences, instead of being suppressed (Figure 2C, dotted region). Hence, the winner-take-all behaviour in the EI circuit became less clear. Instead, there were now three attractor states with a strong central choice-neural attractor (Figure 3A, EI, left inset) (Wong & Wang, 2006). This mixed information led to lack of input separability for the AS group (Figure 3A, right) while receiving overall higher input to urge it to respond faster. Subsequently, the target selection accuracy was close to random (50%) and the go-RT was faster with a narrow range of ∼400-445 ms across coherence levels (Figure 3B, AS, bottom inset).

**Figure 3.**
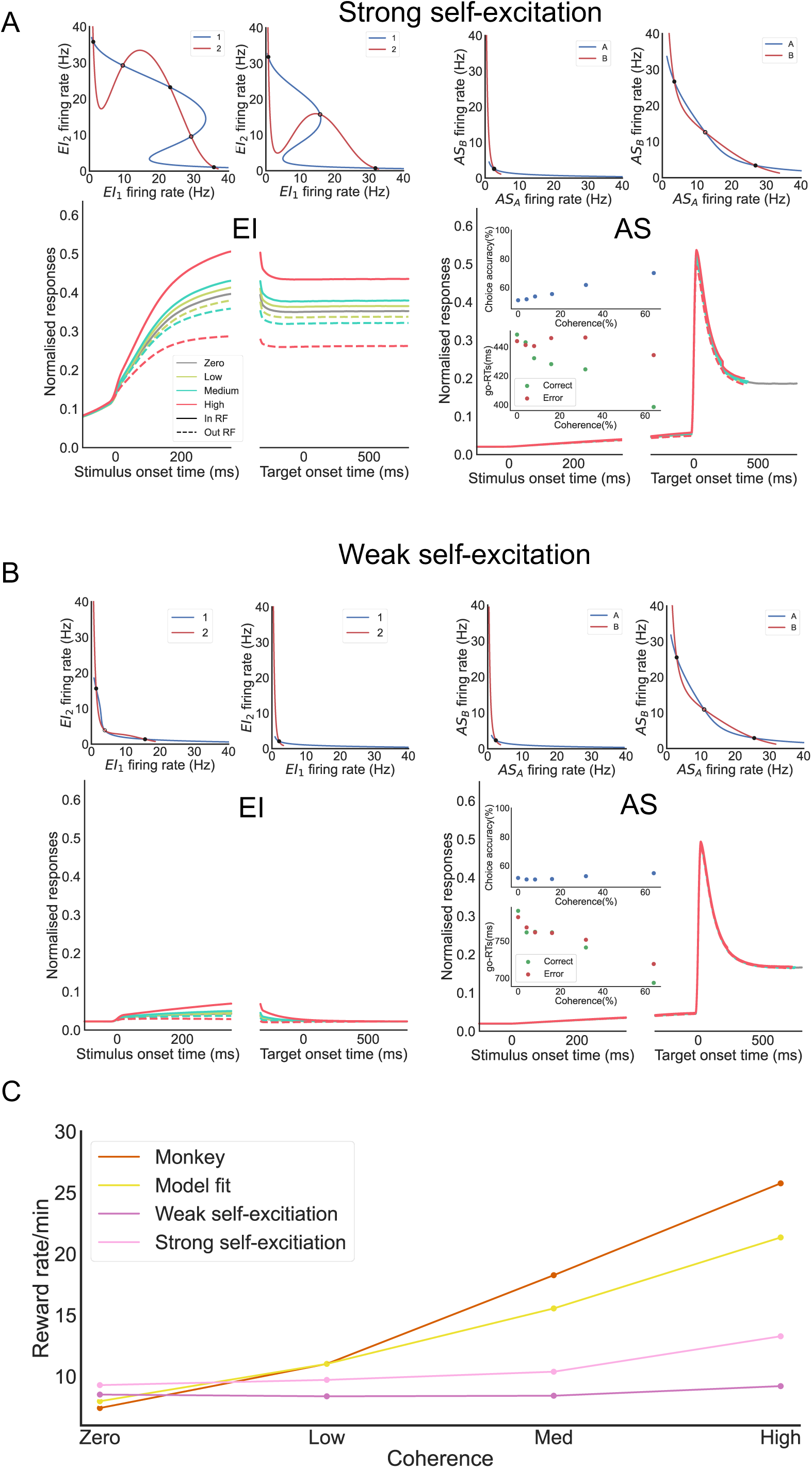
Model predicts optimal recurrent excitatory strength. **(A)** With strong self-excitation in EI neuronal populations, there is higher tendency to be activated but separability between competing neuronal populations is compromised carrying across the delay period (left). This leads to lower separability of inputs to AS neuronal populations (right), resulting in faster go-RT but poorer accuracy. **(B)** With weak self-excitation, EI activities are lower and decays quickly (left), leading to less information into AS (right), and hence slower go-RT and poorer accuracy. (A-B) Labels as in Figure 2. Phase planes with 0% coherence. **(C)** Reward rate of monkey AN (red) and fitted model (yellow) generally outperform model with stronger (light purple) or weaker (dark purple) EI self-excitation. At very low coherences, strong self-excitation marginally attains the highest reward rate.

With weaker EI’s self-excitatory connectivity (𝑆_𝐸,𝐽_ =0.8), the EI activities were generally lower and unable to sustain in the absence of the motion stimulus during delay period (Figure 3B, left). This led to lower and less input separability for the AS group (Figure 3B, right), which again caused random choices but slower go-RT with a narrow range of ∼700-800 ms across coherence levels. This suggests that the fitted model, and hence, monkey AN could be performing near the optimal regime in the processing of decision related information, memory storage and conversion to action selection.

To show this, we incorporated monkey AN’s go-RT, delay period duration (333 ms) and time delay (2500 ms) after making error choices as well as correct choices (500 ms), and computed its overall reward rate across the trials (i.e. the percentage of correct choices divided by the averaged overall time in a trial) (see Methods, and Niyogi & Wong-Lin (2013)). Figure 3C demonstrates that the fitted model, and hence monkey AN, generally attained larger reward rates than with stronger or weaker EI self-excitation. Thus, intermediate recurrent excitatory strength facilitated optimal memory sampling for decisions in this task.

Interestingly, this optimal strategy did not hold for very low motion coherences, i.e. highly difficult tasks such as with ambiguous stimulus (Figure 3C, yellow and red). In such cases, particularly in blocked trials with similar coherence levels, the best strategy, with the highest reward rate, was to have EI be endowed with strong self-excitation (Figure 3C, light purple). Intuitively, this is conceivable as when a task becomes too difficult (e.g. stimulus close to being ambiguous), decisions should be made rapidly given that they are mostly at chance level anyway.

So far, the model accounts for observations from experimental task that was designed to investigate perceptual decision abstracted from effector. In comparison, many studies on perceptual decisions make use of reaction time task in which the stimulus is present until motor response (Roitman & Shadlen, 2002). Hence, we next tested the model’s flexibility to adapt to such task with only reconfiguration of the timing of the stimulus inputs, but no change in the model parameters (Supplementary Table S1).

### Mean-field model accounts for LIP’s distinct neuronal clusters and predicts decision readout mechanism in a different decision task

Recent studies on LIP neurons of non-human primates have made use of multi-neuron/unit recording, revealing interesting low-dimensional neuronal activity dynamics in perceptual decision-making and action selection (e.g. Bollimunta & Ditterich, 2012; Steinemann et al., 2024; Zylberberg & Shadlen, 2025). In particular, heterogeneous LIP neuronal groups had been identified in reaction-time task, in which some LIP neuronal groups exhibited non-stereotypical ramping-to-threshold activity patterns (Zylberberg & Shadlen, 2025). More specifically, Zylberberg and Shadlen (2025) showed that although the averaged LIP neuronal activity might resemble stereotypical ramping-to-threshold pattern (Figure 4A, left), when decomposed, there were three distinct clusters of LIP neurons. One cluster (Cluster 1) encoded sensory evidence strength akin to encoding decision confidence (Figure 4A, middle), a second cluster (Cluster 2) encoded choice information with graded ramping-to-threshold pattern (Figure 4A, right), and a third group encoded decision uncertainty (not shown, but see Cluster 3 in Zylberberg and Shadlen (2025)). Such distributed neural encoding is also in line with our previous modelling work across cortical layers based on human neuroimaging data (Asadpour & Wong-Lin, 2025).

**Figure 4.**
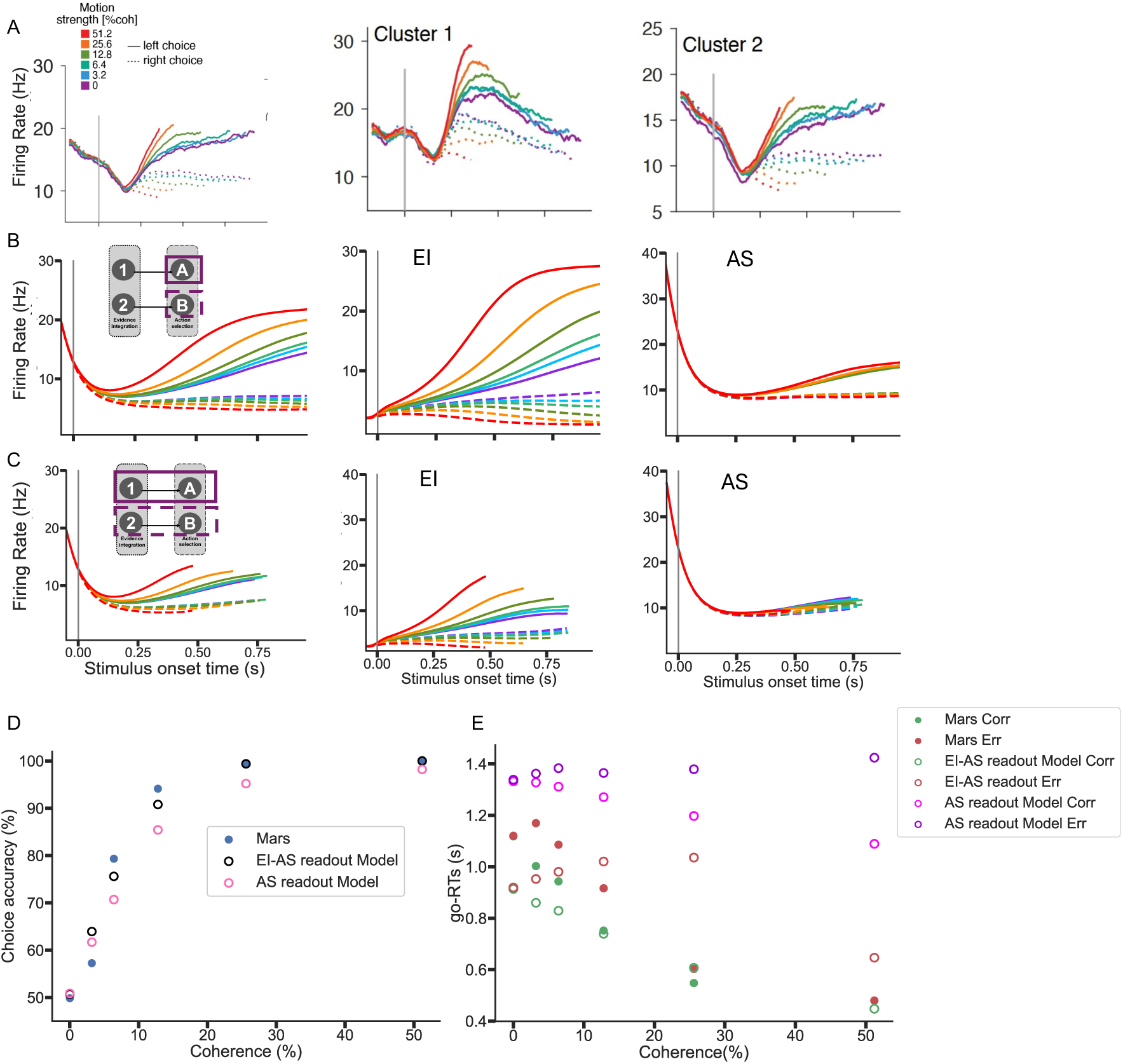
Sensory evidence encoding neural population in motion discrimination reaction time task. **(A)** Simultaneous recording of multiple LIP neurons in non-human primate unveils distinct clusters of neurons (Steinemann et al., 2024; Zylberberg & Shadlen, 2025) Left: Firing rate activities averaged over all recorded LIP neurons; middle: activity of a cluster of neurons that encode task difficulty; right: another neuronal cluster with stereotypical ramping-to-threshold activities encoding choice. Adapted from Zylberberg & Shadlen (2025, Figure 1F and Figure 4C) **(B-C)** Model recapitulates neural and behavioural data of Zylberberg & Shadlen (2025) Same model as in Figures 1-3, but with choice target input preceding motion stimulus input as in (A) (Steinemann et al., 2024). Left: Activities averaged over all model neuronal populations; middle (right): only EI’s (AS’s) averaged activities. Colour labelling as in (A). (D-E) Comparing model psychometrics readout from AS group (B) or averaged of EI and AS groups (C) with monkey Mars in Steinemann et al. (2024).

As decision uncertainty has previously been modelled using minimal mean-field approach (Atiya et al., 2019, 2020, 2021), we shall focus on sensory evidence and choice encoding neurons to demonstrate how our parsimonious EI-AS model can instantiate such computation. In this decision task by monkey Mars (Steinemann et al., 2024), the choice targets were presented before the random-dot motion stimulus onset (Roitman & Shadlen, 2002) while decision confidence or uncertainty was not instructed or evaluated. Hence, the same EI-AS model in Figures 1-3 now had the choice target input before motion stimulus input, as in Wong et al. (2007) without changing other model parameters (Methods, and Supplementary Table S1).

We first cropped the neural activities when the AS group reached some prescribed decision threshold (Figure 4B, left). The coherence dependency of the activities averaged over EI and AS neuronal populations generally resembled that of the averaged activities of the LIP neurons (Figure 4A, left). We then replotted the model activities of the EI and AS groups separately (Figures 4B, middle and right), and found that they resemble qualitatively those of Clusters 1 and 2 of the LIP neurons (Figures 4A, middle and right), respectively. When we considered the decisions to be made collectively by averaging the EI and AS activities, we found that the activity patterns qualitatively followed even more closely those of the LIP neurons (compare Figure 4C with Figure 4A). This suggest that the readout (Supplementary Figure S1) of the collective activities of monkey Mars’ two LIP neuronal clusters, akin to the model’s EI and AS neurons, might be utilised for making perceptual decisions. This was further validated when we plotted the decision accuracy and reaction time (from motion stimulus onset); the psychometrics based on readout from the averaged of EI and AS, as compared to readout from only AS, were more similar to the monkeys’ psychometric data (Figure 4D).

In the above investigations, we have seen how EI neurons in the model can contribute to perceptual decision reporting via action selection and readout. It would be interesting to know whether EI neurons when perturbed prior to action selection can affect behaviour, a form of cognitive interference. More specifically, how would subsequent AS activities and choice behaviour be affected? We shall next investigate this below via a proposed two-stage version of the perceptual decision-making task.

### Decision-based EI neuronal resetting leads to cognitive interference in two-stage decision task

For the task in Shushruth et al. (2022), if the decision of motion direction is reported prior to the availability of the coloured choice targets’ locations, it is unclear whether and how subsequent response to choose the coloured target will be affected. In other words, suppose we slightly alter the experimental task of Shushruth et al. (2022) to probe the monkeys’ decisions about the motion direction before the monkeys choose the corresponding coloured choice targets (Figure 5A, top), will the accuracies and go-RT for choosing the coloured targets be affected? This is similar to previous studies of two-stage human perceptual decision tasks in which the second decision can be influenced by the first decision (Townsend et al., 2000; Busemeyer & Bruza, 2012; Wang & Busemeyer, 2016; van den Berg et al., 2016; Atiya et al., 2019).

**Figure 5.**
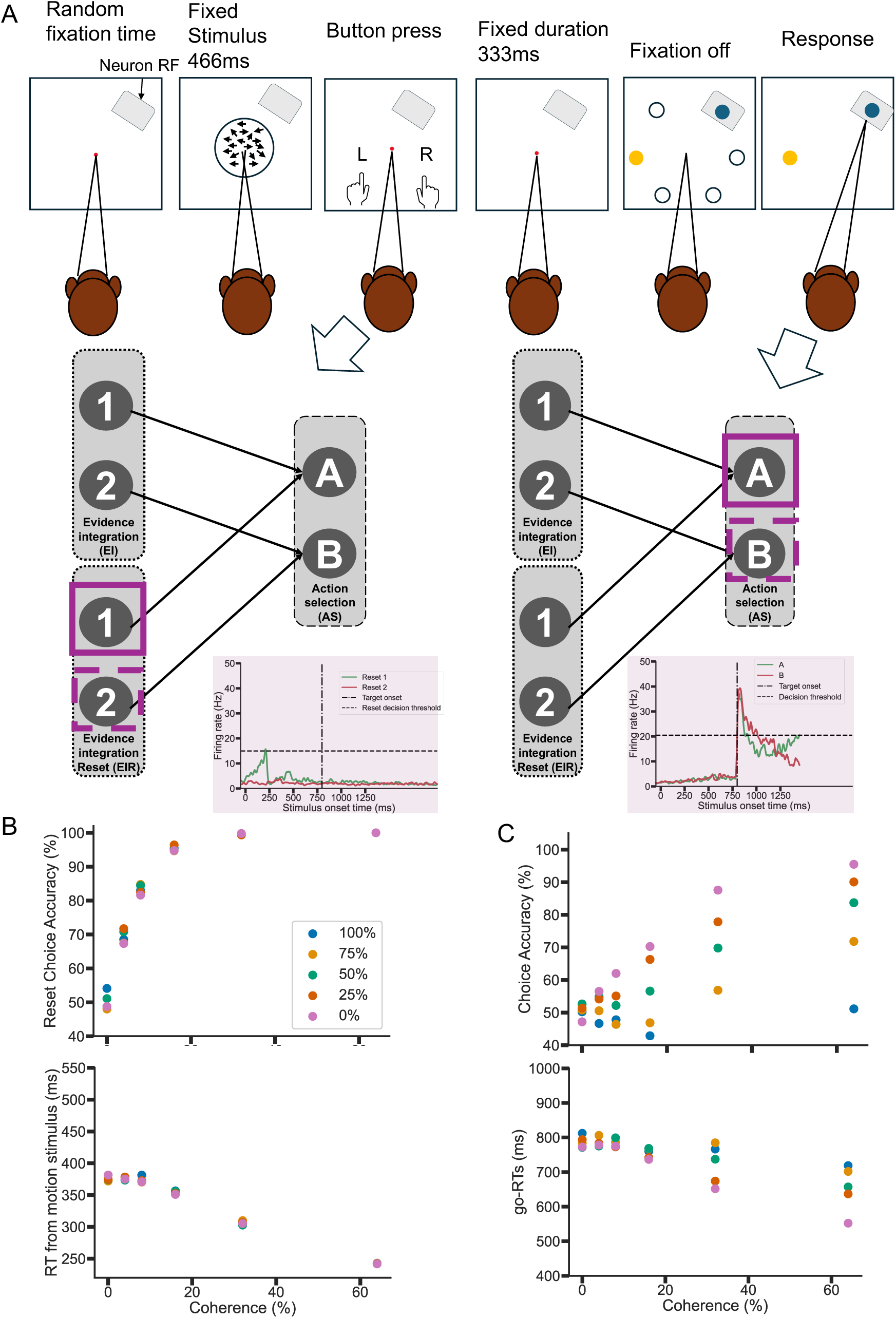
Model with resetting EI neurons predicts cognitive interference in two-stage decision task. **(A)** Top: Two-stage decision task in which the first decision is reported based on motion coherence direction via a different motor modality (e.g. finger pressing) while a second decision is later made as in Shushruth et al. (2022) (Figure 2A). Bottom (left): Model has a fraction of the EI neural group output to be readout to make the first decision (purple boxes), after which their activities are reset to baseline (left inset). Sample activities with 100% of the EI neurons deciding and resetting, and 32% motion coherence. Bottom (right): Model’s AS group makes second decision (via saccade). Purple inset’s horizontal dashed (vertical dashed-dotted) line: decision threshold (target onset time). **(B-C)** Model predicts little effect on first decision as fraction of resetting EI neurons increases (B); there are more interference (detrimental) effects of first decision on second decision’s accuracy than go-RT (C), consistent with lower AS activity separability (right inset in A).

With regards to the above model, we have seen how the EI neuronal group contributes to sensory evidence integration and forming and storing a decision. Hence, a trivial answer would be that there are some downstream neurons that readout the EI’s activities, and the AS neural activities should reflect similar coherence-dependence information from the EI neurons as shown in Figure 2. However, previous studies have demonstrated that when the second decision was associated with the first decision, the second decision’s accuracy decreased – a cognitive interference phenomenon (Townsend et al., 2000; Wang & Busemeyer, 2016). Moreover, our findings above suggest that the readout may involve both EI and AS neurons. Thus, the trivial answer does not satisfy these observations.

Another possibility is that some of the EI neurons may be reset to baseline level upon reaching some decision threshold via inhibitory feedback after motor response (Lo & Wang, 2006; Stine et al., 2023). An experiment to validate this would require some explicit motor response for the first stage. This can be achieved, for instance, using different effector modality such as arm reaching or finger pressing instead of saccadic eye movement in Shushruth et al. (2022) (Figure 5A, top); previous work (e.g. Lafuente et al., 2015) has shown than LIP neurons robustly encode decision variable regardless of reaches or saccades.

We systematically investigated the above by extending the model into two subgroups of EI neurons (Figure 5A, bottom). One (EI) subgroup integrated sensory evidence and stored the decision as before, but another (EIR) subgroup additionally reset upon reaching some decision (Methods; Supplementary Table S1) before coloured target onset (Figure 5A, bottom left inset). As such resetting mechanism led to less coherence-based information being fed to AS, there was less clear coherence-dependent separability in the AS activities (Figure 5A, bottom right inset). Indeed, as the percentage of resetting EI neurons increased (Methods), the first decision’s accuracy and reaction time clearly remained largely unaffected (Figure 5B) but the second decision’s accuracy decreased, as observed in previous studies (Townsend et al., 2000; Busemeyer & Bruza, 2012; Wang & Busemeyer, 2016). Further, the model predicted go-RT to increase (Figure 5C).

## Discussion

We have proposed a neurobiological and minimal mean-field neural circuit model of evidence integration and action selection for gated and flexible perceptual decision-making. The model, inspired by LIP neuronal responses in perceptual decision-making, is input-gated without the need for re-training for different tasks. In particular, the interplay between external inputs and the network’s nonlinear dynamics allows rapid reconfiguring mechanisms of gating and adaptability without re-learning. In particular, we have shown that the network model utilises nonlinear dynamics for decision information to be integrated, cached and acted across different perceptual decision-making tasks. The model additionally provides the simplest account of neural computation operating in curved neural manifolds in perceptual decision-making (Figure 1C) (Thura et al., 2022; Langdon et al., 2023; Perich et al., 2025).

This minimal mean-field model offers a plausible explanation to the discovery of sensory evidence encoding cluster in Zylberberg & Shadlen (2025) – our model, with the same model parameter values as in the Shushruth et al (2022) task, suggests that it might exist to allow rapid system reconfiguration when the task changes, for example, from reaction time perceptual decision tasks to tasks with fixed delays or abstracted from effectors (Roitman & Shadlen, 2002; Shushruth et al., 2022; Gherman et al., 2024; Xie et al., 2024). This postulation could be validated in future experiments. Further, this reinforces our prediction of the existence of such (yet to be observed) neuronal population in Shushruth et al. (2022), which may reside within area LIP (Zylberberg & Shadlen, 2025) or in other brain regions (Charlton & Goris, 2024; Hasnain et al., 2025; Xie et al., 2024; Asadpour & Wong-Lin, 2025). If confirmed, such neuronal population, which encodes decision signals abstracted from specific actions, can not only be used for computing decision confidence (Zylberberg & Shadlen, 2025), but also flexibly and overtly report decisions in various motor modalities such as through saccade, vocalisation, reaching, pointing and finger pressing, or even covert internal mentalisation. As an aside, the readout of both EI and AS neurons provided better fit to the observed neuronal and behavioural data of Zylberberg & Shadlen (2025), and with faster RT, than readout of only AS neurons. Improving such readout strategy is reminiscent of learning algorithms in more abstract reservoir-like networks (Sussillo & Barak, 2013; Song et al., 2017; Xu et al., 2024; Langdon & Engel, 2025; Genkin et al., 2025).

When the model was fitted to the data of Shushruth et al. (2022), we found that even though the task required binary discrimination (leftward-vs-rightward motion perception) or action (choosing blue-vs-yellow target), the neural decision information storage during the delay period was not binary but graded (Figure 2C, left), unlike previous classical modelling (Wang, 2002; Wong and Wang, 2006). In other words, information leakage was needed such that over many trials, the information on average became graded. This in turn led to the prediction of other experimental observations (Figures 4B and 5C). The decay-induced graded decision information encoding has also been previously modelled when accounting for electroencephalography (EEG) activity across temporal gaps (Azizi & Ebrahimpour, 2023), wherein the model was also based on the mean-field model of Wong & Wang (2006). An advantage of such model is its robustness in encoding graded information over time without the need to perfectly fine-tune model parameters into line attractors for storing continuous information (Seung, 2003; Machens et al., 2005; Mante et al., 2013).

Working memory of the committed decision could be attributed to the recurrent excitation in neural networks (Wong & Wang, 2006; Wang, 2021). We showed that the fitted model attained higher reward rate performance than with stronger or weaker EI’s self-excitatory connectivity. This inferred that the monkey (AN) in Shushruth et al. (2022) might have learned to perform the perceptual decision task optimally or close to optimal. Although we only varied the EI’s self-excitation to avoid introducing additional model parameter, we expect similar results with input-driven variation in EI, as previously shown (Wong & Wang, 2006). With such implementation, we expect similar results even in trials with mixed task difficulty, in which some stimulus-induced input can rapidly trigger memory storage capability. The modelling here enables investigation for neural computations of a wider class of decisions particularly memory-based sequential sampling (Shadlen & Shohamy, 2016).

Our parsimonious mean-field model was developed with the motivation to identify the simplest possible neurobiological model that explains and predicts a diverse set of neural and behavioural data in flexible perceptual decision-making. For explainability purposes, we did not specifically model multiple spatial choice target locations as in the task of Shushruth et al. (2022). This would require more complex ring attractor or neural field models (e.g. F. Liu & Wang, 2008; Furman & Wang, 2008; Esnaola-Acebes et al., 2022; Monsalve-Mercado et al., 2025), which may exhibit qualitatively similar decision separability dynamics (You & Wang, 2013). Extensions of the minimal mean-field model could be built for more complex cognition (e.g. Loeffler et al., 2023) or accounting for more complex neuronal selectivity (e.g. Tye et al., 2024), while offering more transparent insights through reverse engineering approach. Reward-based adaptive learning (Soltani & Izquierdo, 2019) could also be incorporated into the minimal mean-field model, for instance to perform reversal learning of stimulus-action maps via EI-AS connectivity.

An intriguing finding in our study was that in a task with two related sequential decisions, the first reported decision could affect subsequent decision within the same trial (Figure 5C). Such cognitive interference, a violation of classical probability theory, was previously accounted for by quantum decision theory (Buseymeyer & Bruza, 2012). Specifically, quantum interference provides a mathematically elegant account of decision interference effects (Townsend et al., 2000; Busemeyer et al., 2009; Wang & Busemeyer, 2016). Our model offers an alternative, neural mechanistic explanation of this interference, which not only recapitulated the second decision’s lower accuracy but also predicted its slower decision-based responses – the EI’s activity resetting upon the first made decision led to less information made available for the second decision. This could re-open research avenues to investigate the neural mechanisms underlying various cognitive interference effects (Sarason et al., 1996).

In summary, we have proposed a biologically plausible minimal mean-field neural circuit model that is inspired by parietal physiology and provides theoretical insights and explanations in flexible perceptual decision dynamics. The model provides a foundational theoretical framework to investigate neural mechanisms of diverse task contexts, including perceptual, memory-based, and abstract decision-making.

## Data availability

Source code, generated data and analyses will be made available upon publication.

## Acknowledgments

We thank Abdoreza Asadpour, Michael Shadlen, Redmond O’Connell, Simon Kelly and Elisabeth Parés-Pujolràs for useful discussions, and Michael Shadlen for providing helpful feedback to the manuscript. This work was supported by HSC R&D (STL/5540/19) and MRC (MC_OC_20020) (AA, KW-L). BL was supported by Ulster University via Northern Ireland Department for the Economy (DfE). We were grateful for access to the Tier 2 High Performance Computing resources provided by the Northern Ireland High Performance Computing (NI-HPC) facility funded by the UK Engineering and Physical Sciences Research Council (EPSRC), Grant No. EP/T022175/1.

