## Supplemental file for "Minimal mean-field gated parietal circuit model for flexible perceptual decisions"

### Supplementary Information

Brendan Lenfesty[1], Amin Azimi[1], Saugat Bhattacharyya[1], S. Shushruth[2], KongFatt Wong-Lin[1,*]

[1]Intelligent Systems Research Centre, School of Computing, Engineering and Intelligent Systems, Ulster University, Magee campus, Derry~Londonderry, Northern Ireland, UK

[2]Department of Neuroscience, Center for Neural Basis of Cognition, University of Pittsburgh, Pittsburgh, USA

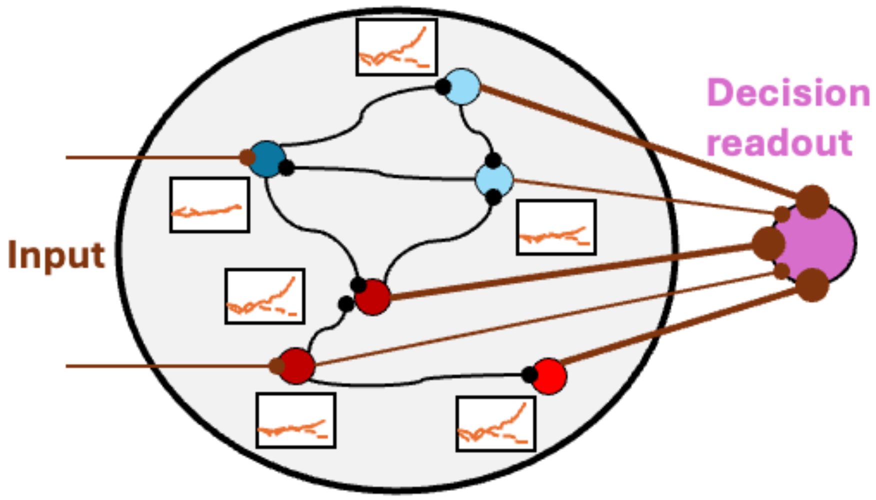


Supplementary Figure S1. Schematic illustrating neurons exhibiting different levels of decision-related activity separability (orange traces) and, consequently, contributing differently, through appropriate connectivity weightage (brown lines), to the decision readout. Assumption: neurons with a stronger relationship to the decision contribute more to the decision process.

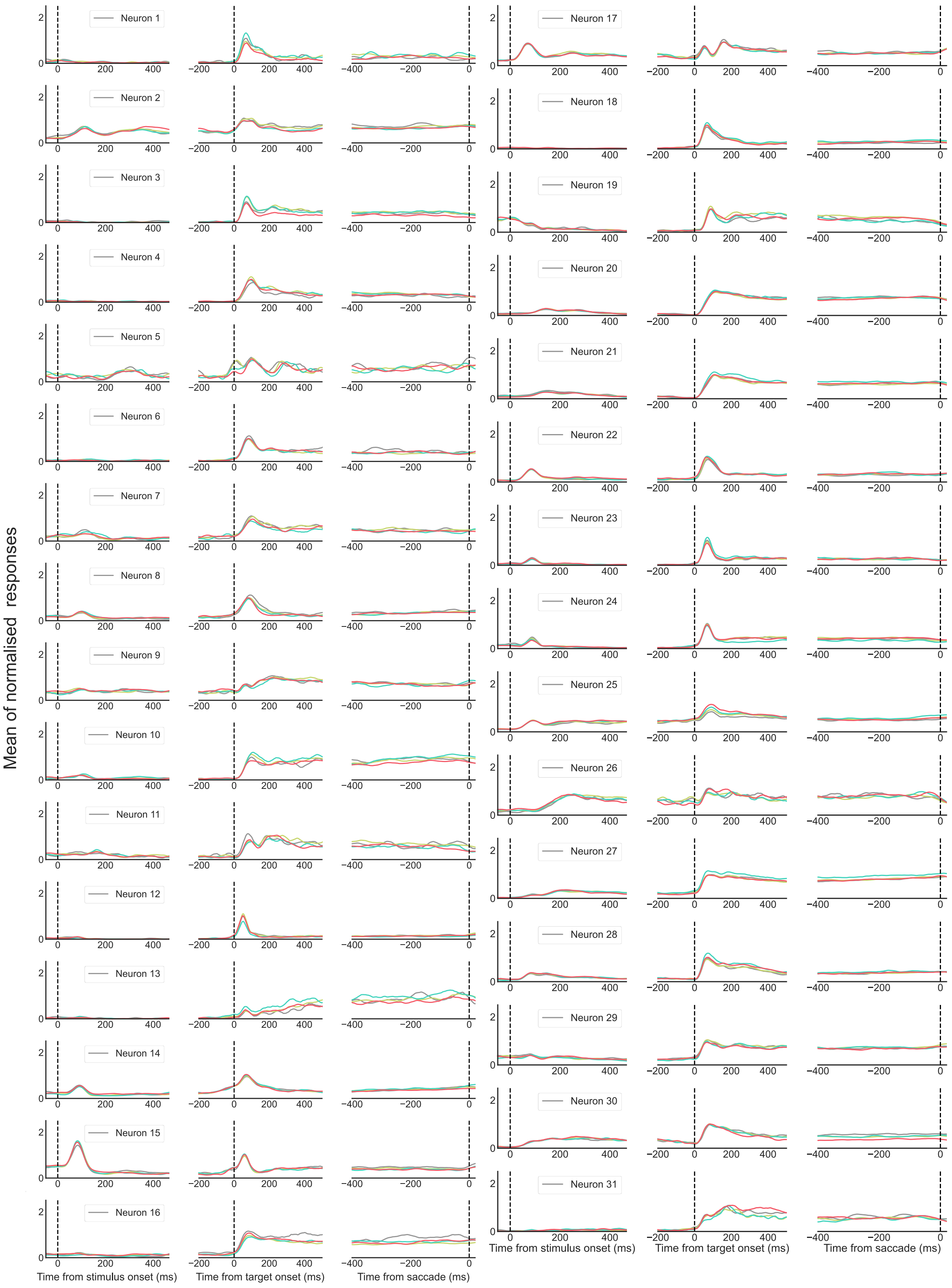


Supplementary Figure S2. Trial-averaged LIP neuronal firing rates for all sessions across all epochs of the go-task experiment for Monkey SM. Labels as in Figure 2 in main manuscript.

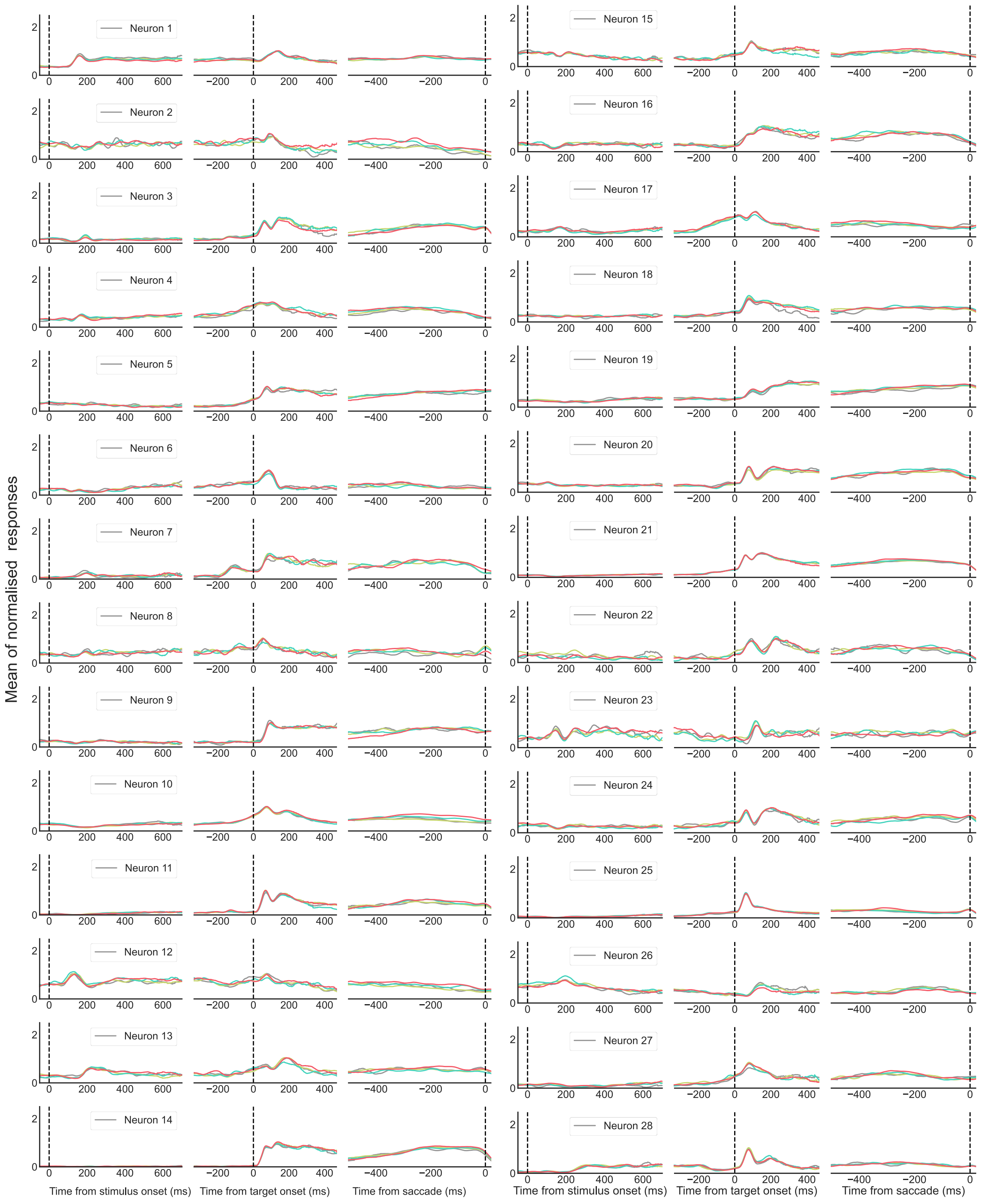


Supplementary Figure S3. Trial-averaged LIP neuronal firing rates for all sessions across all epochs of the wait-task experiment for Monkey AN.

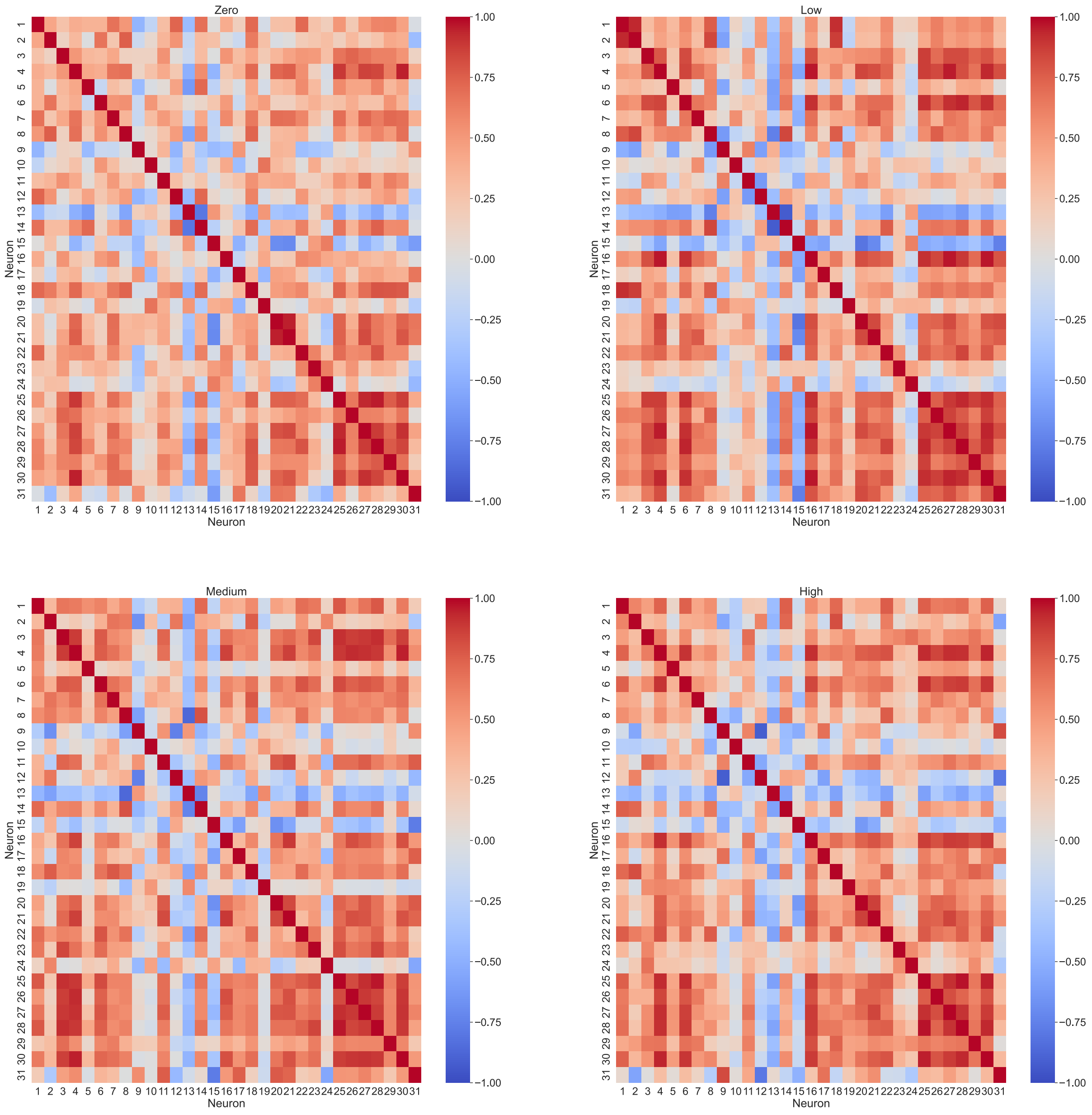


Supplementary Figure S4. Correlation coefficient matrices for neurons/sessions for Monkey SM based on motion coherence levels.

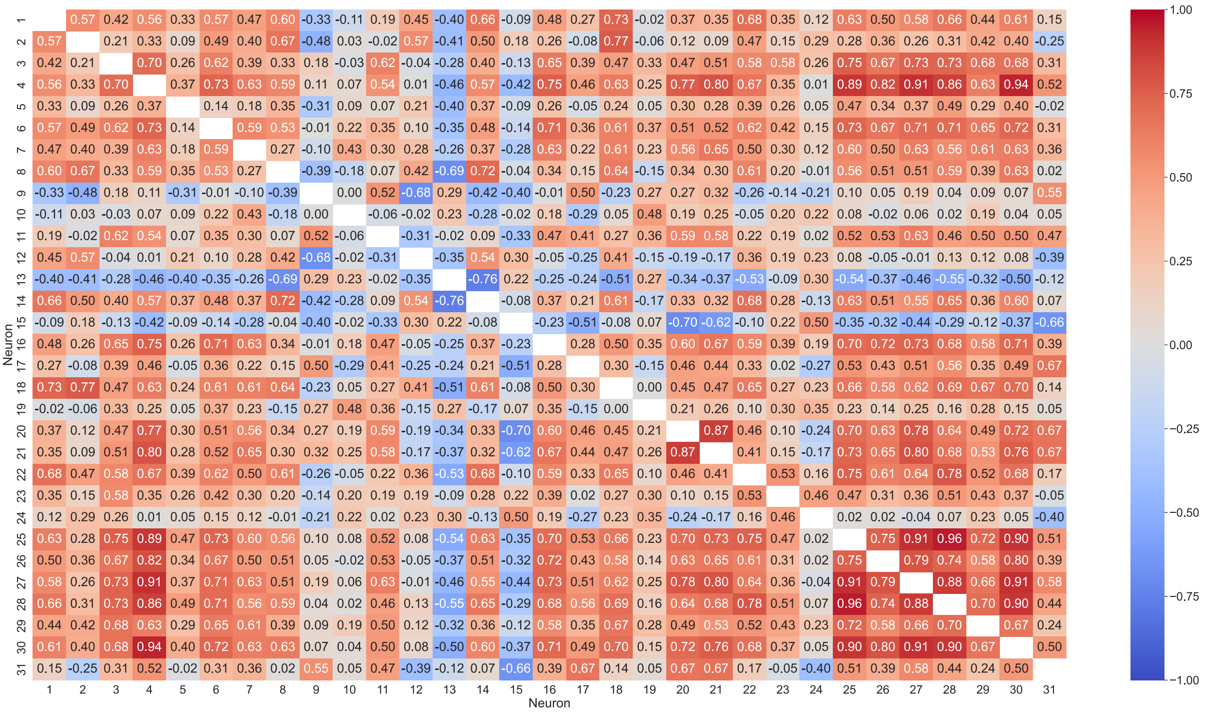


Supplementary Figure S5. Correlation coefficient matrix for neurons/sessions for Monkey SM averaged across motion coherence levels.

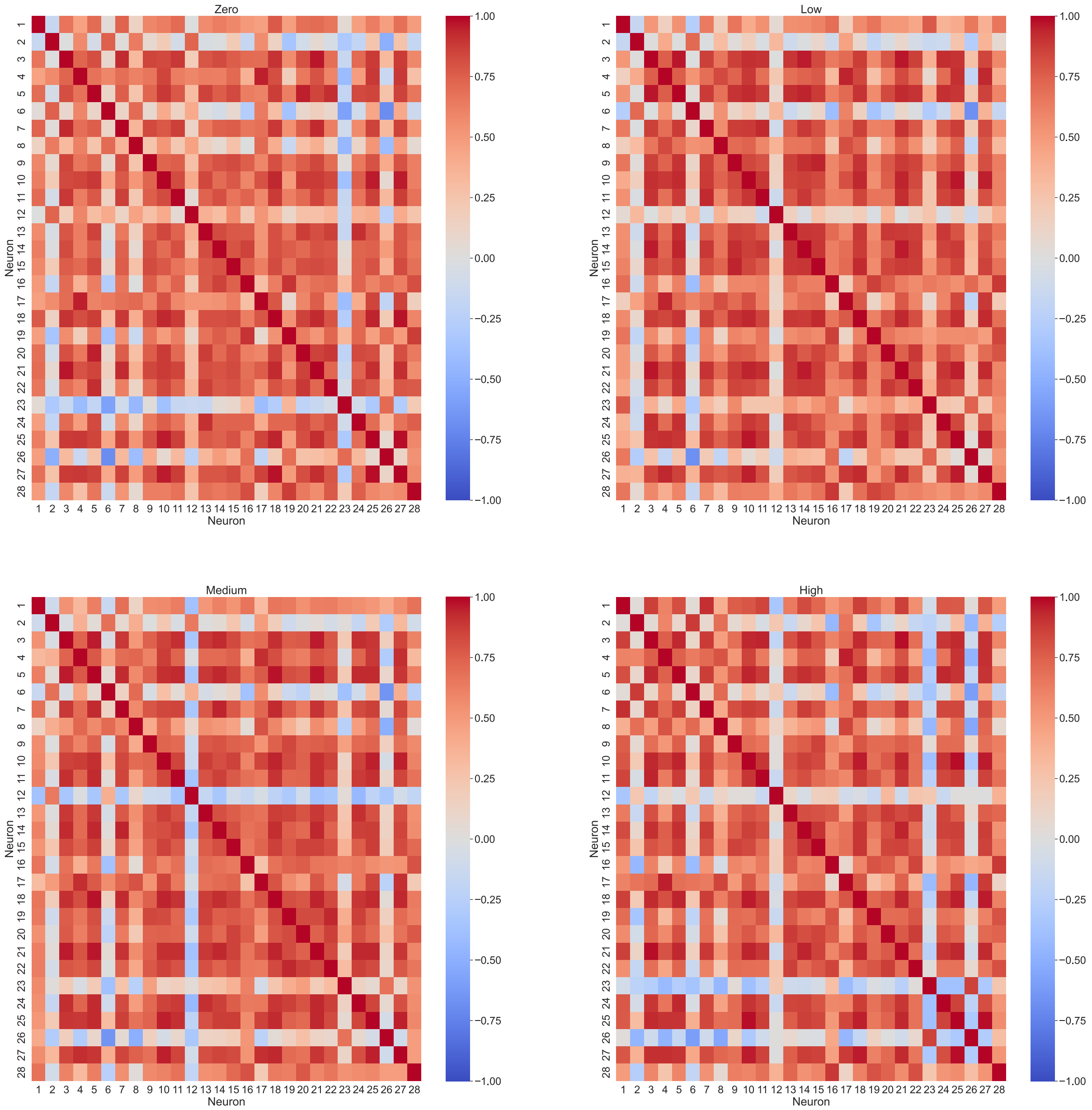


Supplementary Figure S6. Correlation coefficient matrices for neurons/sessions for Monkey AN based on motion coherence levels.

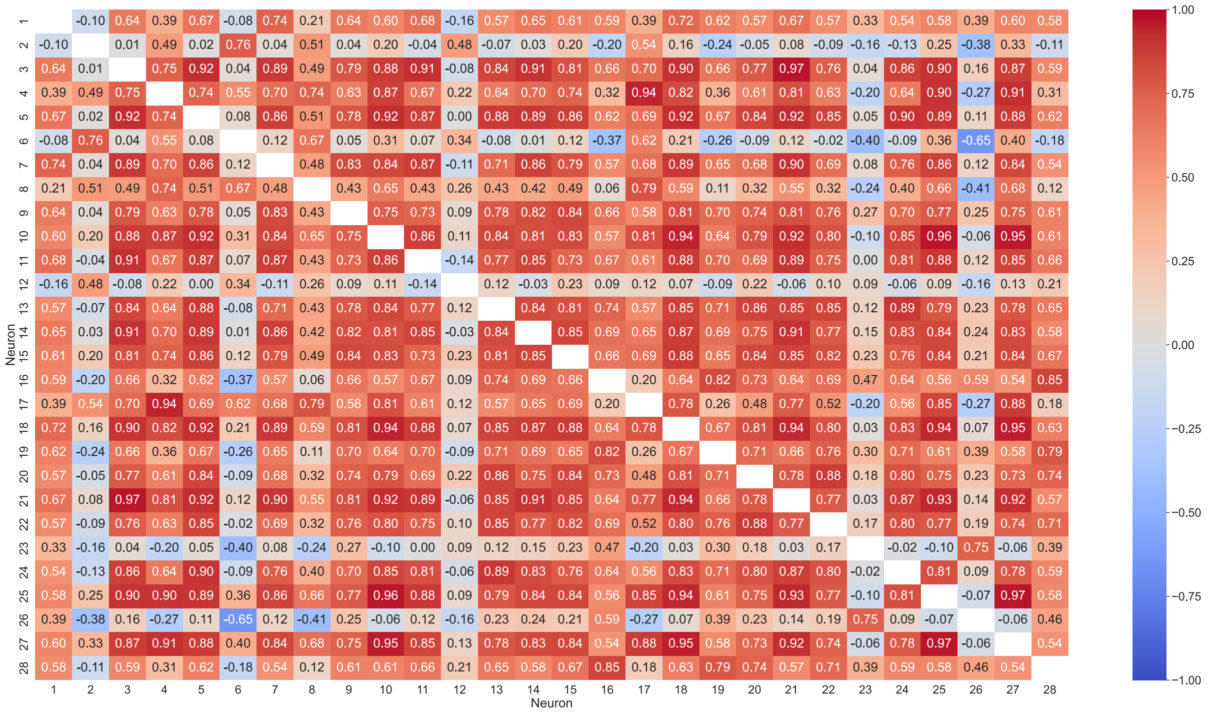


Supplementary Figure S7. Correlation coefficient matrix for neurons/sessions for Monkey AN averaged across motion coherence levels.

| Model parameters | Description | Monkey AN fit (Shushruth et al., 2022)) | Monkey SM fit Shushruth et al., 2022)) | Monkey Mars (Steinmann et al., 2023; Zylberberg & Shadlen, 2025) | Cognitive interference |
|---|---|---|---|---|---|
| $a$ | Neuronal input-output function gain ($(\mathrm{VnC})^{-1}$) | 270 | | | |
| $b$ | Neuronal input-output function bias (Hz) | 108 | | | |
| $d$ | Neuronal input-output function curvature (s) | 0.1540 | | | |
| $I_0$ | Biased current (nA) | 0.3255 | | | |
| $\mu_0$ | Baseline stimulus input (Hz) | 26.5 | | | |
| $J_{ext}$ | External input coupling strength ($\mathrm{nA} \cdot \mathrm{Hz}^{-1}$) | 0.00052 | | | |
| $\tau_{NMDA}$ | NMDA-mediated synaptic decay time constant (ms) for excitatory recurrent coupling | 100 | | | |
| $\tau_{AMPA}$ | AMPA-mediated synaptic decay time constant (ms) for noise generation | 2 | | | |
| $J_{EE,EI}$ | Self-excitation coupling weight in EI (nA) | 0.255 | | | |
| $J_{EE,AS}$ | Self-excitation coupling weight in AS (nA) | 0.17 | | | |
| $J_{inh,EI}$ | Effective inhibitory coupling strength in EI (nA) | -0.0397 | | | |
| $J_{inh,AS}$ | Effective inhibitory coupling strength in AS (nA) | -0.0475 | | | |
| $g_{gain,0}$ | Gain amplitude in AS | 0.790 | | | |
| $\tau_{gain}$ | Gain decay time constant in AS (ms) | 333 | 200 | | |
| $\tau_{ad}$ | Choice target's adaptive time constant in AS (ms) | 60 | | | |
| $g_{coh}$ | Motion stimulus' gain factor | 0.805 | | | |
| $S_{E,J}$ | Multiplicative factor for self-excitation coupling weight in EI | 0.975 | | | |
| $w_{EI-AS}$ | EI-to-AS coupling strength (nA) | 0.0055 | | | |
| $\zeta$ | Stimulus-dependent multiplicative noise | 0.00096 | | | |
| $\sigma$ | Noise amplitude (nA) | 0.0216 | | | |
| | Decision threshold (Hz) | 20.5 | | | 15 (1st decision); 20.5 (2nd decision) |

Supplementary Table S1. Model parameters and their values for the different monkeys and tasks. To allow direct transfer of model parameters from one task to another, all parameter values were maintained to be the same, except for the gain decay time constant for monkey SM and the first decision threshold for the cognitive interference task. Delay period for replicating monkey AN in abstract decision-making task and the cognitive interference task was 333 ms; it was 200 ms to replicate monkey SM while not applicable for monkey Mars. Reaction time was computed from time to crossing decision threshold plus 225 ms non-decision latency.
